# Gut metabolite IPA alleviates white matter post-ICH injury by enhancing myelin debris phagocytosis via *Stap1* inhibition

**DOI:** 10.1101/2025.11.19.689382

**Authors:** Meichang Peng, Meiqin Zeng, Hao Tian, Chen He, Lu Zhang, Zhanyi Zhao, Yunhao Luo, Xin Li, Fangbo Xia, Siyi Liu, Xiao Cheng, Haitao Sun

## Abstract

**Background:** Intracerebral hemorrhage (ICH) causes neurological dysfunction and white matter injury (WMI) characterized by myelin loss, axonal injury and myelin debris accumulation. Microglia-mediated debris clearance is critical for WMI repair. The microbiota-gut-brain axis plays an essential role in the central nervous system diseases, one of the ways in which gut microbiota affects brain is via producing metabolites. Indole-3-proprionic acid (IPA), a tryptophan-derived metabolite that mainly produced by *Clostridium sporogenes*, exhibits anti-inflammatory and neuroprotective properties. However, its effect on ICH remains unclear. This study aims to investigate the IPA level after ICH and the therapeutic effects of IPA on neurological deficits and WMI, as well as the potential mechanisms underlying IPA-mediated neuroprotection.

**Methods:** An ICH model was established using C57BL/6 mice, which then received intragastric IPA (20 mg/kg/day). Fecal abundance of IPA-related genes and IPA levels in feces and plasma were measured by qPCR and UPLC-MS/MS. Behavioral tests, qPCR, and immunofluorescence staining were used to assess neurological function, myelin integrity, and axonal injury. *In vitro*, BV2 microglia, with or without *Stap1* knockdown, were co-cultured with myelin debris to assess IPA’s effects on phagocytosis. Additionally, targeted plasma IPA profiling was performed in 30 ICH patients and 30 matched healthy controls.

**Results:** The relative abundance of IPA production-related genes and IPA levels in feces and plasma were significantly decreased after ICH and remained low into the chronical phase. After IPA administration, the neurological deficits, myelin loss and axonal injury of mice with ICH were significantly improved. *In vitro*, IPA increased BV2 microglia myelin debris phagocytosis by inhibiting *Stap1* expression. IPA levels were significantly reduced in ICH patients, consistent with ICH mouse model.

**Conclusions:** Our findings demonstrated that the gut microbiota-derived metabolite IPA could facilitate neurological deficits recovery and alleviate WMI, which could be a promising therapeutic strategy to improve ICH prognosis.

## Introduction

Intracerebral hemorrhage (ICH) is the most devastating subtype of stroke and contributes the highest counts of disability-adjusted life-year across all stroke subtypes ^1,2^. The poor neurological outcomes of ICH patients are largely due to its complex and dynamics pathological process. Firstly, the vessel rupture accompanies with bleeding, and then forms hematoma that causes mechanical compression, leading to gray matter and white matter (WM) direct cell death and structural damage, such as axonal injury and myelin disintegration, resulting in immediate neurological deficits. Although surgery and drugs can be used to alleviate the primary compression, secondary injury after ICH remains to be solved ^3,4^. In the recent years, it was found that myelin debris accumulation would lead to neuroinflammation and inhibit axonal and myelin regeneration ^5,6^. Myelin debris can be cleaned by phagocytes through phagocytosis and degradation, which may promote neurological function recovery. Microglia is residential immune cell in the brain as well as a types of ‘professional’ phagocytes. Modulating myelin debris clearance by microglia is beneficial for remyelination and neurological function recovery ^7^.

Indole-3 propionic acid (IPA) is a metabolite specific to the gut microbiota, which is mainly produced by *Clostridium sporogenes* (*C.sporogenes*) through tryptophan metabolism, and cannot be synthesized by the human body itself ^8^. As a heterocyclic aromatic organic compound with high resonance stability, IPA exhibits anti-inflammatory and antioxidant properties ^9^, and its production relies on phenylacetate dehydratase expressed by the *fldC* gene ^10^. IPA not only exerts local effects in the gut by contributing to the maintenance of intestinal barrier homeostasis—it upregulates ZO-1 to reduce paracellular permeability and increases the secretion of mucins and goblet cell products to strengthen the mucus barrier ^11^. It can also be absorbed by intestinal epithelial cells into the bloodstream ^12^, diffuse through the blood into the cerebrospinal fluid, and cross the blood-brain barrier to reach the brain ^8^. Due to its functions in both the central nervous system and the immune system, IPA has attracted growing research interest. As early as 1999, during the screening of indole compounds with neuroprotective activity against β-amyloid (Aβ), the Y J Chyan team was the first to identify that the endogenous metabolite IPA could significantly inhibit Aβ-mediated neuronal toxicity, specifically, it not only suppresses lipid peroxidation induced by Aβ or DDTC but also exhibits remarkable hydroxyl radical scavenging capacity ^13^. Subsequent studies have further uncovered the multi-target protective mechanisms of IPA. For instance, in a mouse model of sciatic nerve crush injury, IPA promotes axonal regeneration, epidermal innervation, and the recovery of thermal nociceptive sensation by recruiting neutrophils to the dorsal root ganglia ^14^. Studies have shown that IPA not only promotes the restoration of intestinal barrier integrity after ischemic stroke and regulates the Treg/Th17 balance in gut-associated lymphoid tissue, but also improves neurological deficits, alleviates neuroinflammation, and reduces cerebral infarct size ^15^. However, the role of IPA in both the pathogenesis and potential treatment of ICH remains largely undefined, representing a key research gap.

In previous studies, ICH patients accompany with gut microbiota disorder, and gut microbiome diversity decreased, and relative abundant alteration is relative with neurological outcome of ICH patients ^16,17^. A multivariable Mendelian randomization study confirmed associations between 11 gut microbiota-derived metabolites and cerebrovascular diseases, including ischemic stroke, ICH, and subarachnoid hemorrhage. Specifically, higher relative abundances of the genus *Catenibacterium* and the genus *Lachnospiraceae UCG010* were associated with an increased risk of ICH, while a higher relative abundance of the genus *Butyricimonas* was linked to a reduced risk of ICH ^18^. In our previous studies, we have found gut microbiome changed after ICH in mice, and modulated gut microbiome can help neurological deficits recovery ^19,20^. However, the effects of the gut microbiome—specifically, the gut metabolites that might influence post-ICH outcomes were not thoroughly investigated.

The present study focused on the gut metabolite IPA and explored its potential as a therapeutic approach for ICH. The results demonstrated that IPA can promote myelin debris clearance by microglia, facilitate white matter injury recovery, and improve neurological function recovery in mice after ICH. To determine whether our findings in the ICH model are relevant to humans, we examined a cohort of ICH patients. Indeed, the levels of IPA was significantly reduced in ICH patients. Together, this study is the first to demonstrate the role of IPA in myelin debris clearance following ICH. As a readily available, safe, and cost-effective agent, IPA presents a readily translatable therapeutic avenue.

## Methods

Data supporting the findings of this study are available from the corresponding author upon reasonable request. All animals’ experiments were approved by the Ethics Committee of Zhujiang Hospital, Southern Medical University (LAEC-2023-130). All experiments were conducted in accordance with institutional guidelines. Animals were randomly assigned to different groups by coin flipping. The animal allocation was masked to the researchers who established the models, performed the neurobehavioral tests, and analyzed the data. C57BL/6J mice (male, 10-12 weeks of age, 25-30 g) were purchased from Zhuhai BestTest Bio-Tech Co., Ltd.

### Intracerebral hemorrhage mouse model

According to the previous research protocol to establish ICH mouse models ^21^. Briefly, anesthesia was induced with 4% isoflurane and maintained with 2% isoflurane, the mice were placed onto the stereotactic frame. Then, using intracerebral injection of type IV collagenase into the right corpus striatum (coordinates from bregma: 2.0 mm lateral, 0.2 mm anterior, and 3.5 mm ventral), ICH was induced by a slow injection 0.08U type IV collagenase in 0.5 μL saline at a rate of 0.05 μL/min. after the operation, the mice were placed on the heating pad. Once they regained consciousness from anesthesia, they were safely returned to their cages, with access to clean water and food ensured.

### Human cohort

A total of 60 participants were enrolled in this study from Guangdong Traditional Chinese Medicine Hospital, including 30 patients with ICH and 30 healthy controls. The inclusion and exclusion criteria were based on those used in a previous study ^22^. Participants were randomly selected from the hospital population. The study protocol received approval from the Ethics Committee of Guangdong Provincial Hospital of Chinese Medicine (Approval number: YE2022-369-01), and written informed consent was obtained from all participants before their inclusion.

### IPA administration

As previous described ^14^, IPA (57400, Sigma) was diluted in sterile PBS at a concentration of 0.5 mg per 200 μL. The mice were treated with 20 mg/kg/day by gavage. Sterile PBS was treated as vehicle.

### Fecal collection

Fecal from mice were collected immediately upon defecation in sterile tubes, and stored at -80℃ until microbiota and metabolomics analysis.

### Plasma collection

Blood was collected from the inferior vena of mice using a syringe, then injected into EDTA tubes, inverted up and down several times, and then centrifuged at 3000 rpm for 20 minutes at 4℃. The supernatant was aspirated as plasm, which was immediately stored at -80℃ until metabolomic analysis.

### IPA quantification

Fecal pellets (2-3 pellets, pre-weighted) and plasma plasma (50 μL) was deproteinized with 3 volumes of ice-cold acetonitrile (150 μL), vortexed for 10 minutes and centrifuged at 14 000 rpm, 4 °C for 15 minutes. The supernatant was dried under a gentle nitrogen stream, and the residue was sequentially treated with 50 μL derivatization reagent, 50 μL HATU and 50 μL TEA, followed by vortexed at 37 °C for 15 minutes. After a second centrifugation at 14 000 rpm for 15 minutes, the final supernatant was transferred to an autosampler vial and analyzed by UPLC-MS/MS for the quantification of IPA as previously described ^23^.

### 16s rRNA sequencing

16s rRNA sequencing was performed by Novogene Co., Ltd. The specific microbiota analysis was conducted followed by previously described ^19^.

### Neurobehavior test

All mice were undergoing one-week training period to reach the baseline level before the formal tests. Two different assessments were involved: open field test and gait analysis. For the open field test, each mouse was allowed to move freely in a 50 cm×50 cm×50 cm test box, and its activity was recorded for 5 minutes. EthoVision XT 15 was used to analyze the mouse’s movement speed and distance. For the gait analysis, each mouse was allowed to walk naturally in the passage, and the software automatically collected the information of the footprints left by the mouse when it passed through the passage. CatWalk XT 10.7 was used to analyze the mouse’s footprints. To minimize the interference with mice, we only assessed their neurological function at the pre-operation stage, as well as on the day 3 and day 14 post-ICH.

### Immunofluorescence staining analysis

The brain tissue of mice was collected, fixed with 4 % paraformaldehyde at 4℃ for 24-48 hours, then paraffin-embedded samples were prepared followed by serial sectioning, and 4 μm continuous coronal slices were prepare for immunofluorescence. Briefly, after antigen retrieval and blocking using 10% sheep serum, the slices were incubated with the primary antibodies at 4 °C for overnight. Then, the slices were incubated with fluorescein-conjugated antibodies at room temperature for 1 hour. The nuclei were stained with DAPI (Solarbio). The images were taken using an inverted fluorescence microscope (Ti2-E, NIKON). Fluorescence intensity was quantified by using Image-J software. The primary antibodies used included Anti-NeuN (1:500, Abcam), Anti-MBP (1:400, Novus Biologicals), Anti-NF200 (1:400, Sigma-Aldrich), Anti-APP (1:200, Proteintech), and Anti-SMI32 (1:100, BioLegend). The fluorescein-conjugated antibodies used included Alexa Fluor®488 Anti-Rabbit (1:400, Abcam), Alexa Fluor®488 Anti-Rat (1:400, Abcam), Alexa Fluor®594 Anti-Mouse (1:400, Abcam), Fluor®647 Anti-Rabbit (1:400, Abcam).

### Bulk RNA sequencing

Ipsilateral striatum of brain tissues were collected from mice that had IPA (20 mg/kg/day) or sterile PBS administration after ICH and stored at -80℃. These samples were sent to Shanghai Biotree Biomedical Technology Co., Ltd. for transcriptome sequencing and result analysis.

### Real-time quantitative PCR (RT-qPCR)

Ipsilateral striatum of brain tissues were collected and homogenized in Trizol reagent , and total RNAs concentration were determined by Multiskan sky (*Thermo Scientific*). Then each 1 μg RNA was reverse transcribed to cDNA using *Evo M-MLV* reverse transcription assay kit (AG11705). The resulting cDNA was subjected to real-time PCR with gene-specific primers in the presence of SYBR Green pre-mix qPCR kit (AG11739) using the PCR system, as described previously^20^. β-actin (*Actb*) was used as an internal standard control and relative gene levels were detected by the 2^-△△Ct^ method. The primers sequences are shown in Additional file 1: Table 1.

### Clostridium sporogenes and fldC gene detection

The genomic DNA was extracted from the mice fecal samples using the *SteadyPure* fecal DNA extraction kit (AG21036) and subjected to RT-qPCR for the detection. The primers sequences are shown in Additional file 1: Table 1.

### Cell culture and transfection

BV2 microglia cells were donated by Dr. Qing Jia from Zhujiang hospital, Southern Medical University. For constructing knockdown of Stap1, BV2 cells were transfected with siRNA (HanYi Biotech). BV2 cells transfected with si-*Stap1* RNA were designated si-*Stap1*, and BV2 cells transfected with empty vectors were designated Control (The validity of the transfection sequence is shown in the Supplementary Fig 1). Cells were grown in DMEM (Gibco) with 10 % fetal bovine serum and Penicillin-Streptomycin Solution (Gibco). Cells were cultured in a 37℃ incubator with a 5% CO_2_ atmosphere.

### Myelin isolation and purification

Myelin was isolated from C57BL/6 mice (8-10 weeks old) brain according to a described protocol ^24^. Briefly, the brain tissues were placed onto a dish on ice and cut into small pieces. Then the tissues were homogenized with a tissue homogenizer in an ice-cold 0.32 M sucrose solution. The brain homogenate in 0.32 M sucrose solution was gently transferred onto the top of 0.83 M sucrose solution in a 50-mL polypropylene centrifuge tube and centrifuged at 20000 rpm for 65 minutes at 4℃. Myelin debris was carefully collected from the interface between the sucrose densities, and then resuspended with Tris-HCl and centrifuged at 20000 rpm for 65 minutes at 4℃. The pellet was resuspended with Tris-HCl and centrifuged at 20000 rpm for 45 minutes at 4℃. Next, the pellet was resuspended with sterile PBS at 12000 rpm for 10 minutes at 4℃ for obtaining more purified myelin debris. Finally, the myelin debris was resuspended with sterile PBS and stored at -80℃.

### Myelin phagocytosis and degradation assays

For phagocytosis assay, BV2 cells were incubated with myelin debris (50μg/mL) for 2h. After incubation, cells were washed with PBS and fixed in 4% PFA for 15mins. For degradation assay, BV2 cells were incubated with myelin debris (50μg/mL) for 2h and washed with PBS, then cultured in reduced FBS (2 % FBS) media for 48h, then washed with PBS and fixed in 4% PFA. Cell staining myelin debris was tested by immunofluorescence with MBP antibody^25^.

### Statistical analyses

Data are represented as the Mean ± SEM. All statistical analyses were performed using GraphPad Prism 10.0 software. For comparisons between two groups, an unpaired Student’s t-test was applied. Comparisons among more than two groups were performed using one-way ANOVA, followed by Bonferroni’s post hoc test. In addition, non-parametric tests were employed where appropriate. A *P* value of less than 0.05 was considered statistically significant. The significance levels are denoted as follows: NS (no significance); * (*P* < 0.05); ** (*P* < 0.01); *** (*P* < 0.001); **** (*P* < 0.0001).

## Results

### 1. ICH mice exhibit systemic IPA levels reduced

To assess whether ICH alters systemic level of microbiota-derived metabolite IPA, ICH mouse model was established (Fig 2A), and targeted metabolomics analysis (UPLC-MS/MS) was used to evaluate the plasma and faeces of ICH mice. The results revealed a significant and sustained reduction in plasma IPA concentration after ICH compared to pre-ICH (Fig 2B, *P* < 0.0001), persisting for at least 14 days post-ICH (Supplementary Fig 2). Consistent with plasma level, the faecal IPA level was also declined after ICH (Fig 2C, *P* < 0.05).

**Figure 1.**
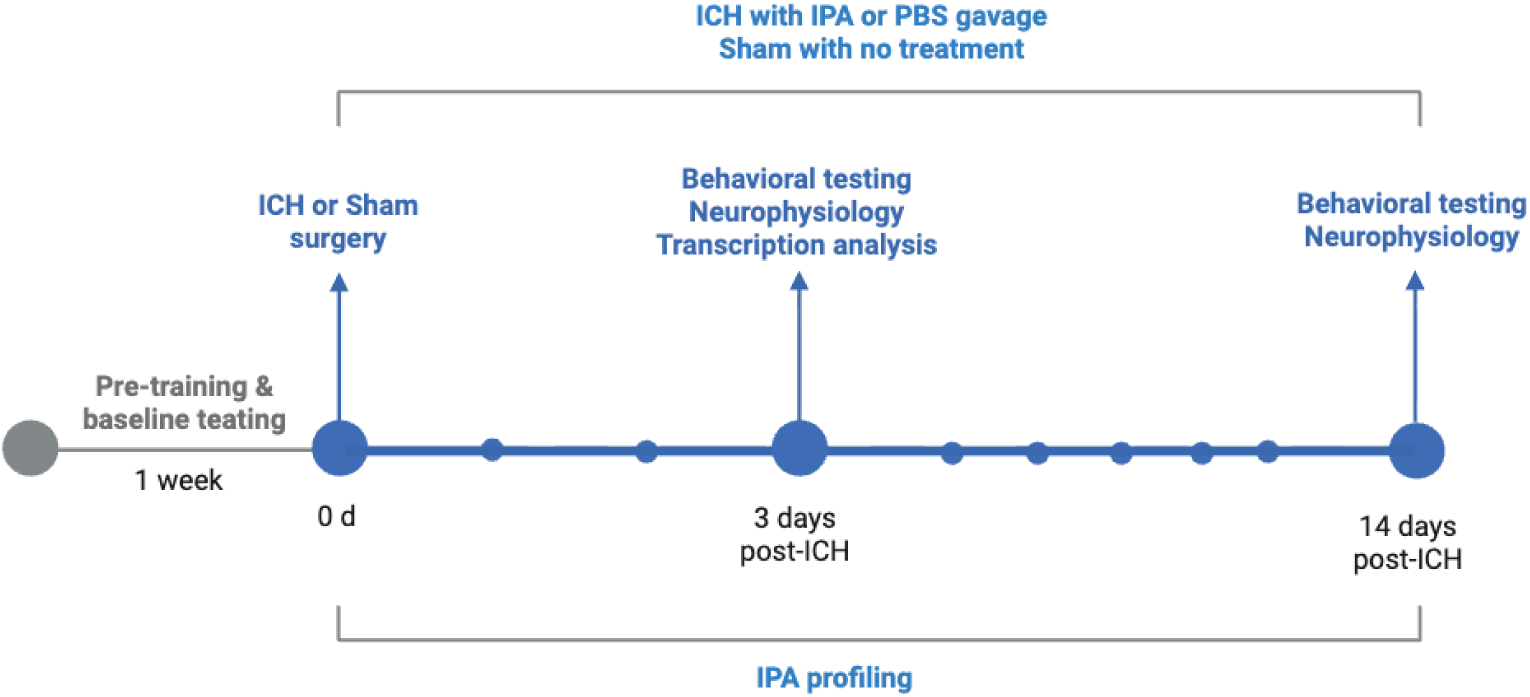
Summary of the experimental procedures, timepoints, and treatment groups in both in vivo.

**Figure 2.**
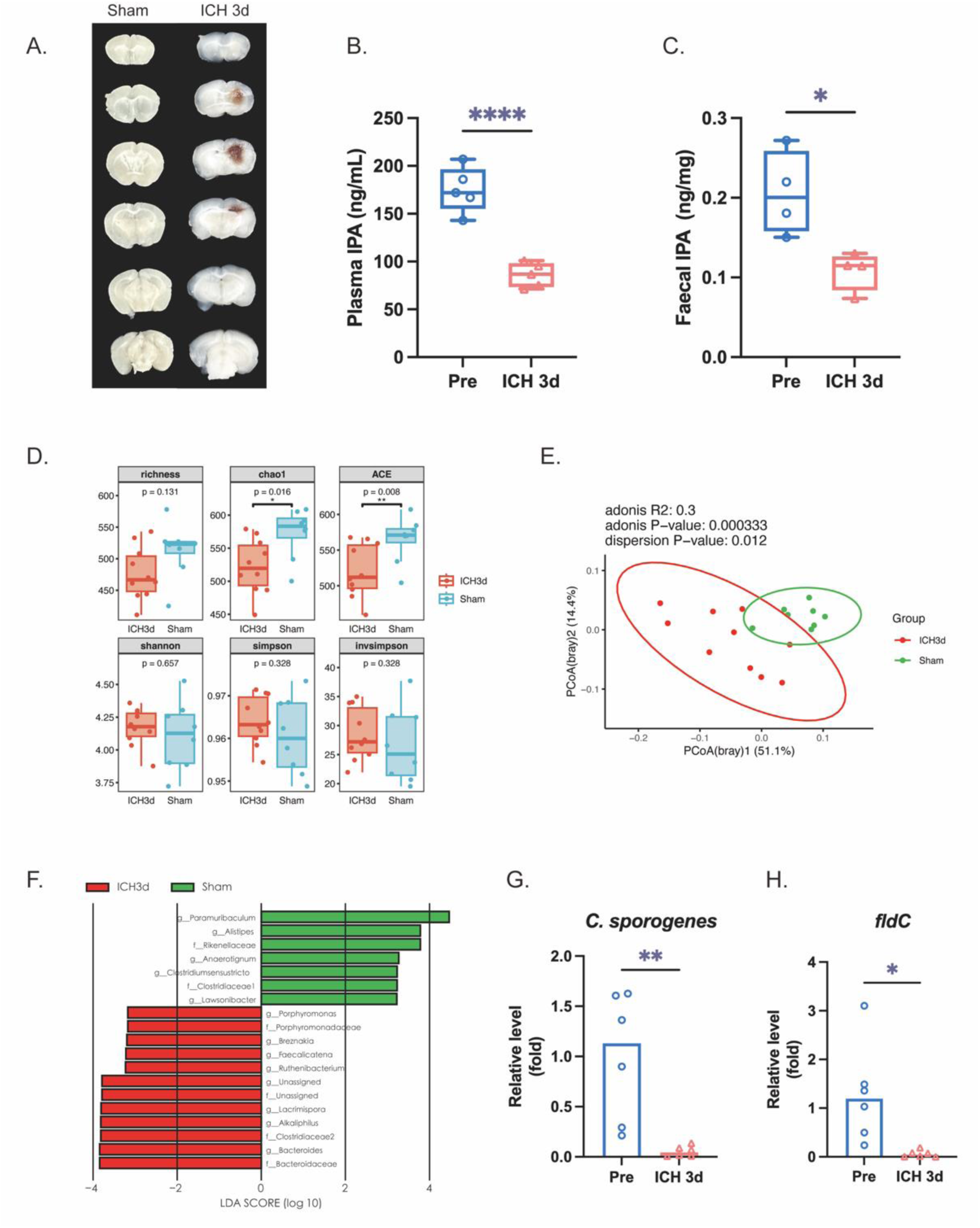
Circulating and fecal IPA levels decrease alongside reduced IPA synthesis in mice after ICH. Plasma and fecal samples were collected from mice before (Pre) and after ICH for analysis by UPLC-MS/MS, qPCR and 16S rRNA gene sequencing. (A) Representative images of brain sections of mice between Sham and ICH groups. (B) Plasma IPA concentrations were quantified by UPLC-MS/MS. Levels were significantly lower at 3 days post-ICH compared to Pre mice (n = 5). (C) Fecal IPA content was quantified by UPLC-MS/MS and was significantly reduced in ICH mice compared to Pre mice (n = 4). (D, E) The α-diversity and β-diversity of gut microbiota in each group (n=8-10). (F) Differentially abundant taxa between Sham and ICH 3d groups identified by LEfSe analysis (LDA score > 2.0) (n=8-10). (G) Relative abundance of *C. sporogenes* in fecal samples, as determined by qPCR, was decreased after ICH (n = 6). (H) Relative expression of the *fldC* gene, encoding a key enzyme for IPA synthesis, in fecal samples was detected by qPCR and found to be decreased after ICH (n = 6). Data are presented as the mean ± SEM. \*\*\*\**P*<0.0001, \*\*\**P*<0.001, \*\**P*<0.01, \**P*<0.05, ns *P*>0.05. D and E by Student t test. C, F and G by one-way ANOVA followed by Turkey’ multiple comparisons test between four groups.

Since faecal IPA level was changed after ICH, we wonder whether the gut microbiota are the reasons. 16s rRNA sequencing was used to profile the gut microbiota composition of ICH and pre-ICH mice. Our results showed that the α-diversity was significantly decreased after ICH (chao1 and ACE index, Fig 2D), and principal coordinate analysis (PCoA) based on Bray-Curtis distance demonstrated a significant difference in the composition of the microbiota after ICH (Fig 2E), which was in line with our previous studies ^19,20^. Additionally, LEfSe analysis identified f_Clostridiaceae1 as a significantly differentially abundant feature between ICH and Sham mice (Fig 2F). Due to 16s rRNA sequencing has limited species-level resolution, it is difficult to accurately quantify changes in the abundance of functional strains. We then detected *C.sporogenes* by qPCR, and our results showed that the relative mRNA level of *C.sporogenes* was decreased after ICH (Fig 2G, *P* < 0.01). qPCR also revealed that *fldC*, a IPA-production key bacterial enzyme coding gene ^10^, was also decreased after ICH (Fig 2H, *P* < 0.05).

Based on the above results, we demonstrated that IPA production by gut microbiota was declined and led to systemic IPA level decreased after ICH.

### 2. IPA treatment promotes neurological deficits recovery after ICH

Since the observed systemic IPA was decreased after ICH, we next investigated whether directly supplementing IPA could affect the progression of neurological damage after ICH. The open-field test and catwalk gait analysis were performed to evaluate neurological function. On the open-field test, IPA treatment enhanced the speed and distance in the field compared to vehicle treatment at day 3 and day 14 (Fig 3A-D). On the catwalk gait analysis, IPA treatment improved the motor coordination at day 3 and day 14 (Fig 3E-H).

**Figure 3.**
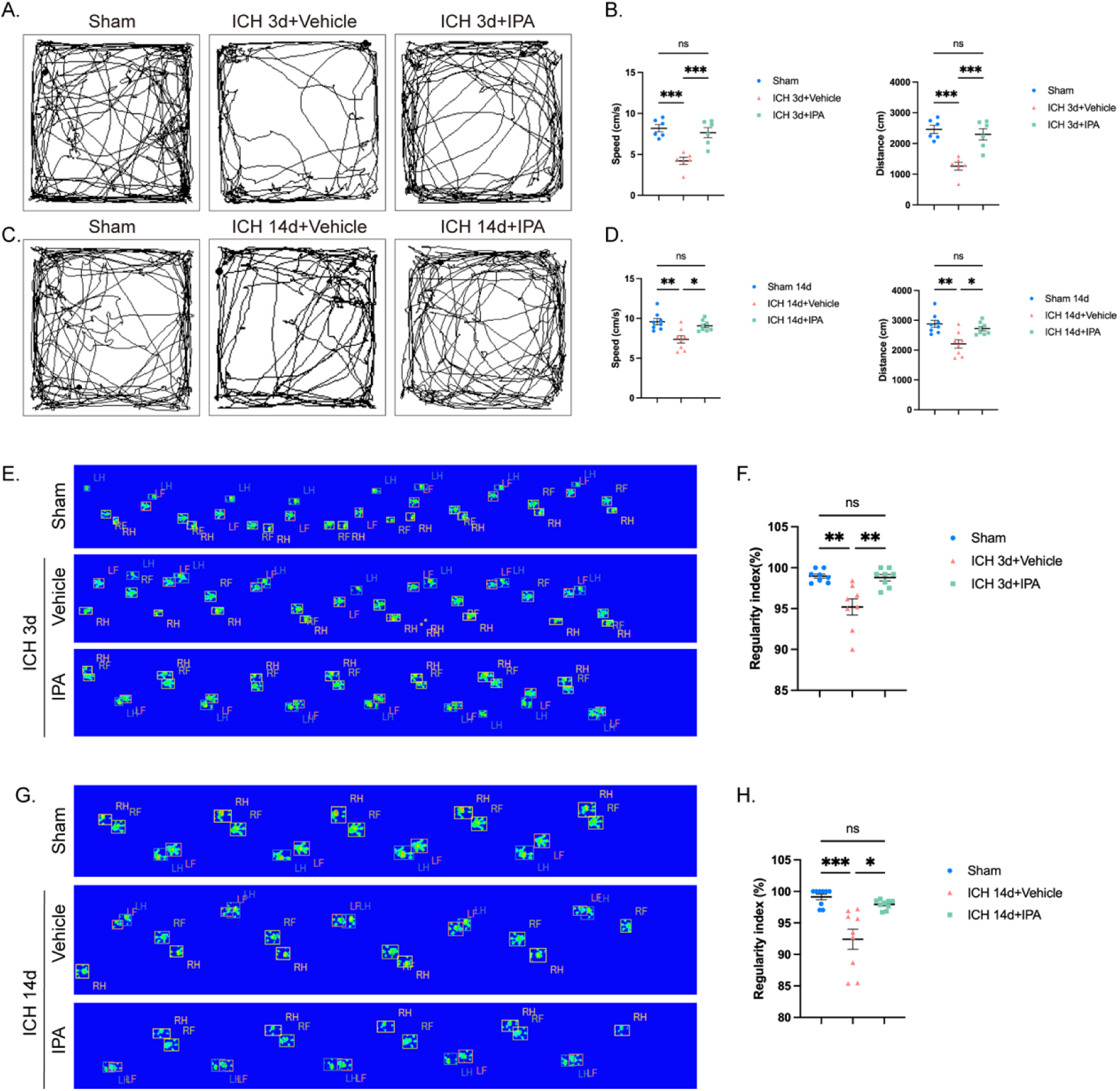
IPA treatment improved neurological functions in mice after ICH. Mice were treated or not treated with 20 mg/kg/day IPA after ICH surgery. (A, C) R Representative trajectories of the open-field test on 3 days and 14 days after ICH. (B, D) Analysis results of speed and distance in the open-field test on 3 days and 14 days after ICH (n = 6-8). (E, G) Representative footprints of the Catwalk gait analysis on 3 days and 14 days after ICH. (F, H) Analysis results of the regularity index in the Catwalk gait analysis on 3 days and 14 days after ICH (n = 8-9). Data are presented as the mean ± SEM. \*\*\**P*<0.001, \*\**P*<0.01, \**P*<0.05, ns *P*>0.05. B, D, F and H by one-way ANOVA followed by Turkey’ multiple comparisons test between three groups.

### 3. IPA treatment preserves white matter integrity by alleviating axonal injury and myelin loss

White matter injury is a critical determinant of long-term neurological deficits after ICH. Unlike neurons, glia cells retain a capacity for repair and regeneration, making them a promising therapeutic target. We therefore assessed whether IPA administration mitigates WMI in the peri-hematoma during both acute and chronic phases after ICH. Our results showed that IPA treatment reduced the APP (a beta precursor amyloid protein that would accumulates abnormally near the injury site ^26^) numbers at 3 days (Fig 4A, E, *P* <0.05) and reduced SMI32 (a marker for axonal injury in chronical phase^27^) intensity at 14 days (Fig 4C, I-J, *P* < 0.01) after ICH, indicated that IPA treatment could alleviate axonal injury. Besides, the mean intensity of MBP after IPA treatment was higher than that of vehicle mice both at 3 days and 14 days after ICH (Fig 4B, F, *P* < 0.05; Fig 4D, K, *P* < 0.01), suggesting that IPA treatment could reduce myelin loss in ICH mice in both acute and chronical phase.

**Figure 4.**
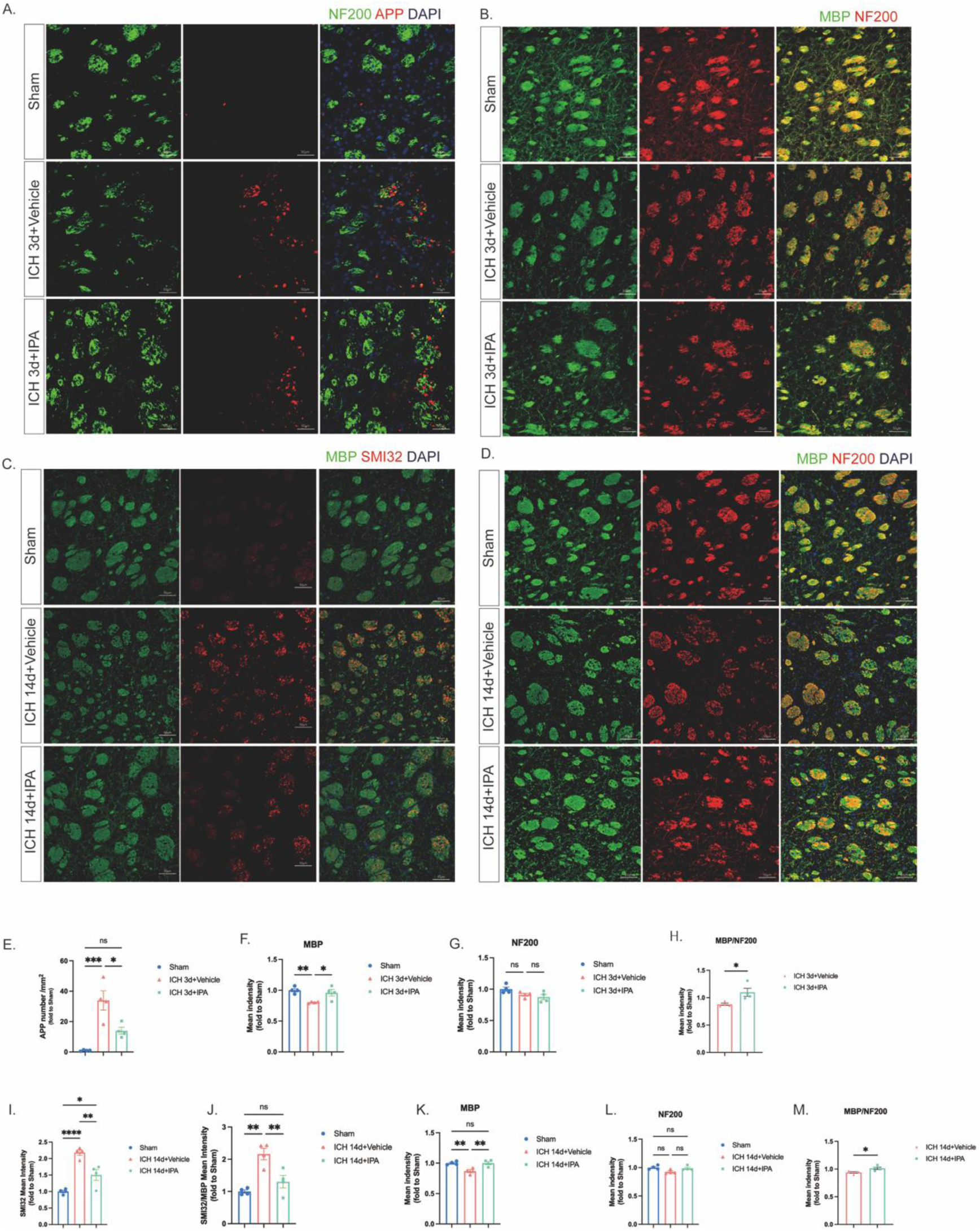
IPA treatment alleviated axonal injury and reduced myelin loss after ICH. (A) Representative immunofluorescence images of NF200 (green), amyloid precursor protein (APP, red) and nucleus (DAPI, blue) in the perihematomal area, showing the axonal injury of the white matter around the hematoma at 3 days after ICH. (B) Representative immunofluorescence images of myelin basic protein (MBP, green) and neurofilament 200 (NF200, red) in the perihematomal area, showing the integrity of the white matter around the hematoma at 3 days after ICH. (C) Representative immunofluorescence images of MBP (green), SMI32 (red) and DAPI (blue) in the perihematomal area, showing the axonal injury of the white matter around the hematoma at 14 days after ICH. (D) Representative immunofluorescence images of MBP (green), NF200 (red) and DAPI (blue) in the perihematomal area, showing the integrity of the white matter around the hematoma at 14 days after ICH. (E) Quantitative analysis results of the number of APP expressions per square millimeter (n = 4). (F, G, H) Semi-quantitative analysis results of the average fluorescence intensity of MBP, NF200, MBP/NF200 (n = 4). (I) The average fluorescence intensity of SMI32 (n = 4). (J) The ratio of the average fluorescence intensity of SMI32/MBP (n = 4). (K, L, M) Semi-quantitative analysis results of the average fluorescence intensity of MBP, NF200, MBP/NF200 (n = 4). Scale bar=50 μm. Data are presented as the mean ± SEM. \*\*\**P*<0.001, \*\**P*<0.01, \**P*<0.05, ns *P*>0.05. H and M by Student t test. E, F, G, I, J, K, L by one-way ANOVA followed by Turkey’ multiple comparisons test between three groups.

### 4. IPA treatment enhances myelin debris clearance of microglia

Myelin debris clearance is crucial for axonal injury recovery; we next want to investigate whether IPA has effect on myelin debris clearance. Our results showed that the mean intensity of Iba-1 and the co-localization coefficient of Iba-1 and MBP in the striatum around hematoma was higher in the IPA treatment group of mice (Fig 5A, B, *P* < 0.05, C, *P* < 0.01), suggesting that IPA may enhance the microglia activity and promote the uptake and clearance of myelin debris by microglia.

**Figure 5.**
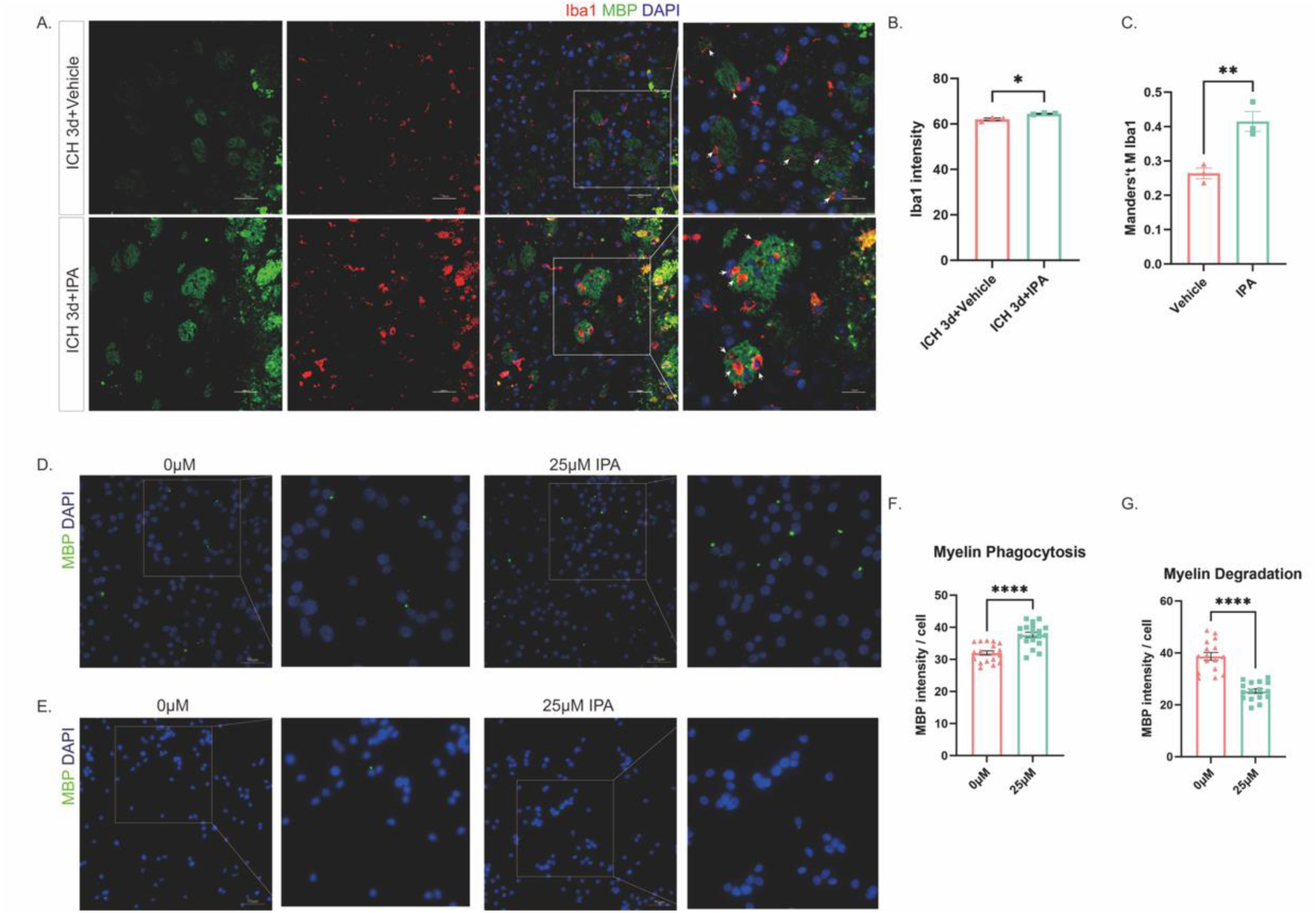
IPA treatment enhanced myelin debris clearance by microglia after ICH. (A) Representative immunofluorescence images of MBP (green), Iba1 (red) and DAPI (blue) in the peri-hematoma area at 3 days after ICH. The white arrows indicate representative microglia phagocytosing myelin debris. (B) Semi-quantitative analysis results of the average fluorescence intensity of Iba1 (n = 3). (C) Analysis results of the co-localization analysis of MBP and Iba1 using the Manders’ coefficient (n = 3). (D) Representative immunofluorescence images of the effect of IPA on the phagocytosis of myelin debris by BV2 microglia. Myelin debris is labeled with MBP (green), and the cell nuclei are labeled with DAPI (blue). (E) Representative immunofluorescence images of the effect of IPA on the degradation of myelin debris by BV2 microglia. Myelin debris is labeled with MBP (green) and the cell nuclei are labeled with DAPI (blue). (F, G) Semi-quantitative analysis results of the average fluorescence intensity of MBP within a single cell in phagocytosis/degradation system. Replicate wells in each group, 5 to 6 visual fields were photographed per well. Scale bar=50 μm. Data are presented as the mean ± SEM. \*\*\*\**P*<0.0001, \*\*\**P*<0.001, \*\**P*<0.01, \**P*<0.05, ns *P*>0.05. B, C, F and G by Student t test.

To further verified that this effect of IPA, we established a co-culture system of BV2 microglia and myelin debris (Fig 6I). By using the mean intensity of MBP to estimate the myelin debris phagocytosis and degradation activity, we found that BV2 cell treated with IPA exhibited significantly higher activity of myelin debris in both phagocytosis and degradation (Fig 5D-G, *P* < 0.0001). These *in vitro* experiments further indicated that IPA promotes the phagocytosis and clearance of myelin debris by BV2 cells. Based on the above results, we speculated that IPA may play an important role in promoting the myelin debris clearance by microglia, thereby reducing white matter damage after ICH.

**Figure 6.**
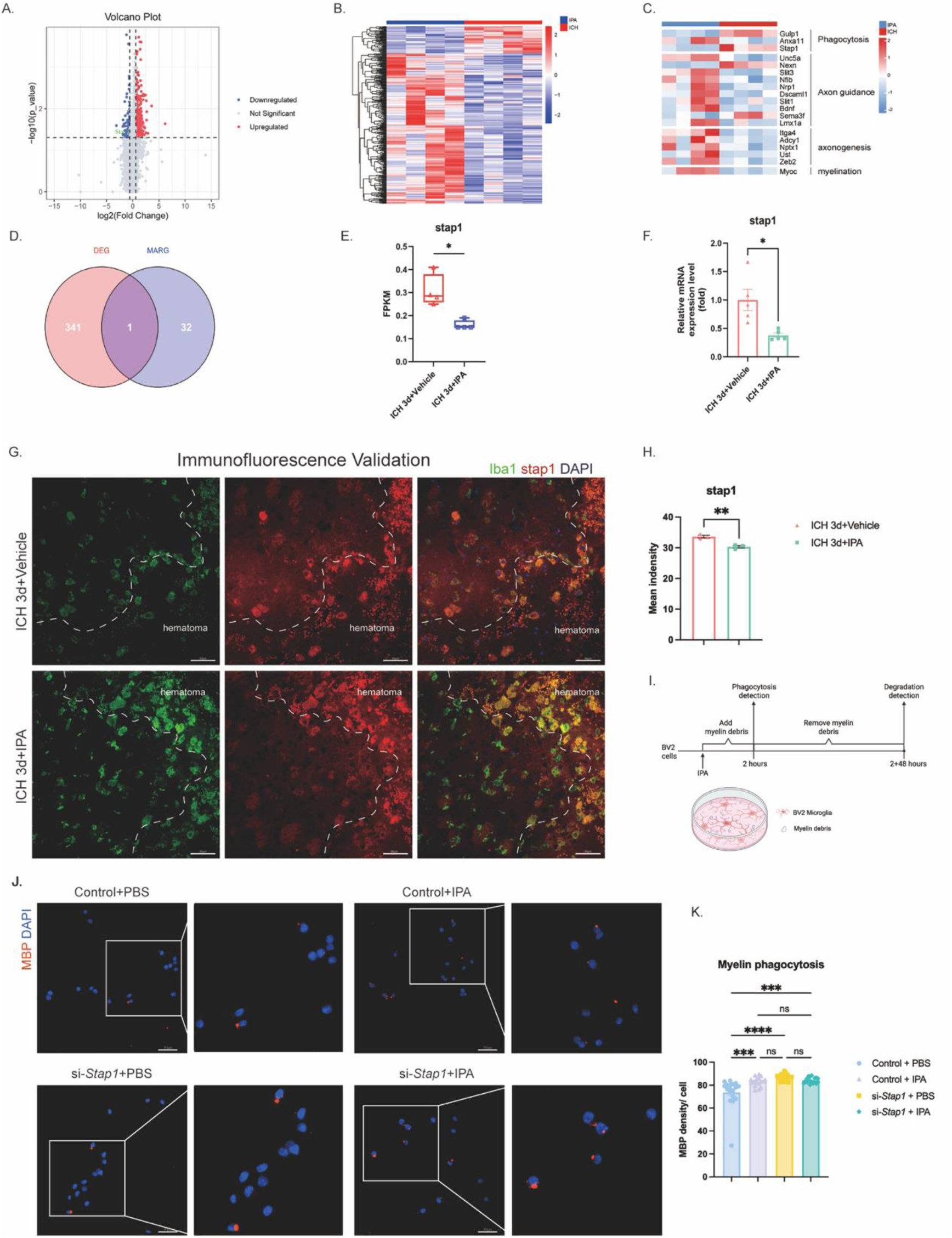
IPA promotes myelin debris phagocytosis of microglia partly through inhibiting *Stap1*. (A) Volcano plot depicting upregulated and downregulated genes and number of genes upregulated and downregulated in ICH + Vehicle mice and ICH + IPA mice at 3 days after ICH (n=4 mice per group; p < 0.05 and log2(fold change) ≥ |0.585|). (B) Heatmap of differential gene transcript in CH + Vehicle mice and ICH + IPA mice at 3 days after ICH (n=4 mice per group; p < 0.05 and log2(fold change) ≥ |0.585|). (C) Heatmap showing the expression patterns of genes of interest in the samples. (D) Venn diagram of the intersection between DEGs and the Gene Ontology database for Microglia Activation Regulation (MARG). (E) Analysis results of the FPKM values of the Stap1. (F) Semi-quantitative analysis results of qPCR for *Stap1* (n = 5). (G) Representative immunofluorescence images of Iba1 (green), STAP1 (red) and DAPI (blue) in the peri-hematoma area at 3 days after ICH. (H) semi-quantitative analysis results of the average fluorescence intensity of STAP1(n = 3). (I) Schematic diagram showing the time course and intervention of BV2 cell and myelin debris co-culture system. (J) Representative immunofluorescence images of the effect of IPA and *Stap1* knockdown on the phagocytosis of myelin debris by BV2 microglia. Myelin debris is labeled with MBP (red) and the cell nuclei are labeled with DAPI (blue). (K) Semi-quantitative analysis results of the average fluorescence intensity of MBP within a single cell in phagocytosis/degradation system. Replicate wells in each group, 5 to 6 visual fields were photographed per well. Scale bar=50 μm. Data are presented as the mean ± SEM. \*\*\*\**P*<0.0001, \*\*\**P*<0.001, \*\**P*<0.01, \**P*<0.05, ns *P*>0.05. E, F, and H by Student t test. K and M by one-way ANOVA followed by Turkey’ multiple comparisons test between four groups.

### 5. IPA enhances microglia phagocytosis by inhibiting *Stap1* expression

To understand how IPA could confer beneficial effects in myelin debris clearance, we performed transcriptomic sequencing on the striatum surrounding the hematoma in the vehicle and IPA treatment in ICH mice. We found that exogenous supplementation of IPA after ICH significantly upregulated 290 genes and significantly downregulated 52 genes (Fig 6A-B). Among them, genes related to phagocytic function including *Gulp1*, *Anxa1*1 and *Stap1* (Fig 6C).

To further investigate the effects on microglia of IPA, we intersected the differentially expressed genes (DEG) with the microglial activation regulation dataset (MARG), and we found that *Stap1* is the intersecting gene (Fig 6D). The result of FPKM indicated that IPA treatment downregulated *Stap1* after ICH (Fig 6E, *P* < 0.05). qPCR and immunofluorescent staining both confirmed that IPA significantly downregulates the mRNA and protein expression of *Stap1* in ICH mice (Fig 6F-H).

For further validation, we constructed *Stap1* downregulated BV2 cell by using si-RNA (si-*Stap1*) and then observed the phagocytosis of myelin debris (Fig 6I). The results showed that IPA treatment significantly enhanced phagocytosis in control cells (Fig 6J-K, Control+PBS vs Control+IPA, *P* < 0.0001). Notably, genetic knockdown of *Stap1* alone similarly enhanced phagocytosis to a level comparable to IPA treatment (Fig 6J-K, Control+PBS vs. si-Stap1+PBS, *P* < 0.0001; Control+IPA vs. si-Stap1+PBS, not significant). Crucially, in Stap1-knockdown BV2 cells, IPA treatment failed to further augment phagocytosis (Fig 6J-K, si-Stap1+PBS vs. si-Stap1+IPA, not significant).

Based on the above results, we are spectacled that IPA enhanced myelin debris clearance partly by inhibiting *Stap1* expression to enhance phagocytosis of microglia.

### 6. IPA levels are reduced in ICH patients

To assess the clinical relevance of IPA, we quantified its plasma IPA levels in a cohort comprising 30 ICH patients and 30 well-matched healthy controls from Guangdong Traditional Chinese Medicine Hospital participants using UPLC-MS/MS analysis. Plasma was collected from patients on the first 24 hours after ICH onset. In this human cohort, IPA levels were significantly decreased in ICH patients compared to healthy controls (Figure 7, *P* <0.05), which is consistent with our findings in the mouse model. Given the severe right-skewed distribution of IPA levels in the human cohort (healthy control), we performed natural logarithmic transformation to ensure the robustness of parametric analyses. Both the log-transformed independent t-test and the non-parametric Mann-Whitney test confirmed a statistically significant difference between the two groups (Supplementary Fig 3).

**Figure 7.**
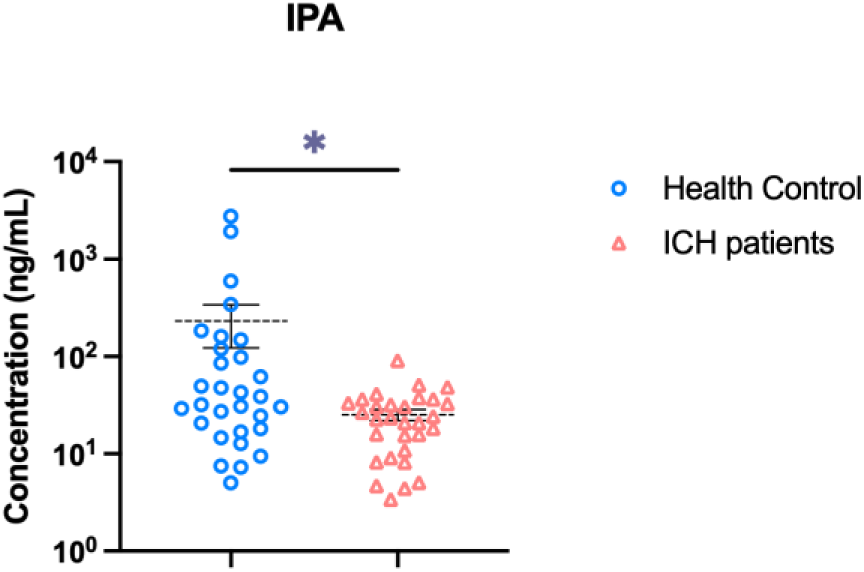
Plasma IPA level was decreased after ICH in human. Plasma was collected from patients 1 day after ICH and from well-matched healthy controls. The level of IPA was quantified using UPLC-MS/MS (n = 30). Data are presented as the mean ± SEM, and the Mann-Whitny test was used for statistical analysis. \**P*<0.05.

## Discussion

To our knowledge, this study provides the first demonstration of the therapeutic effect and underlying mechanism of IPA treatment in ICH mouse model. We have observed 3 key findings: (1) IPA was decreased after ICH in both human and mice, and the reason is the IPA synthesis decreased after ICH; (2) IPA treatment mitigates neurological function and white matter injury; (3) IPA treatment enhances myelin debris clearance of microglia partly by inhibiting *Stap1*. Furthermore, our results align with the established concept that the myelin debris clearance is a critical prerequisite for white matter injury repairment and subsequent neurological recovery (Abstract figure).

The observed declined in systemic IPA level after ICH underscores a critical disruption in the microbiota-gut-brain-axis. Our finding suggested that ICH induces gut microbiota dysbiosis is consistent with previous studies in rodent models ^19,20^ However, beyond general gut microbiota dysbiosis, our results show there exists a deficit in the IPA synthesis pathway after ICH, linking a defined microbial metabolic pathway to ICH pathology.

We also observed that IPA treatment alleviated neurological deficits and WMI. This neuroprotective effect of IPA aligns with beneficial roles in ischemic stroke, autism spectrum disorder and Alzheimer’ s disease models ^15,28–31^. Most ICH patients exhibit WMI. Compared to gray matter injury, WMI demonstrates stronger neuroplasticity and axonal regeneration capacity. Injury to neuronal cell bodies in gray matter can directly lead to cell death, whereas WMI primarily involves axonal damage; if the cell bodies remain viable, cell survival is more easily maintained, facilitating myelin regeneration and axonal regrowth. Therefore, promoting WMI repair could contribute to neurological function recovery after ICH. The process of WMI repair mainly consist of two stages: myelin debris clearance that predominantly mediated by microglia, and myelin regeneration mediated by oligodendrocyte precursor cells (OPCs). After ICH, demyelination and axonal injury in the white matter generate a substantial amount myelin debris. The primary component of myelin debris is lipid ^32^, which can act as an inflammatory stimulant, activating innate immune cells such as microglia and macrophages, thereby promoting the secretion of large quantities of pro-inflammatory factors, including IL-1β and TNF-α ^33^. Furthermore, myelin debris can interfere with the differentiation and maturation of OPCs ^34^. Thus, persistent accumulation of myelin debris in the injured area not only exacerbates neuroinflammation but also inhibits myelin regeneration ^34,35^. Consequently, effectively enhancing the myelin debris clearance is one of the core elements in the repair and regeneration of WMI.

Mechanistically, we identified microglia as the target of IPA in facilitating myelin debris clearance, and IPA partly inhibits microglia *Stap1* to facilitate its ability to phagocytose myelin debris. After injury, microglia are the first cell type to respond, which can be activated within minutes and extend their cytoplasmic processes to the lesion site ^36^. When exposed to the disease-specific microenvironment, microglia enter a disease-associated state and phagocytose invaders ^37^. Subsequently, microglia exhibit various morphologies, such as the amoeboid state (with phagocytic ability), ball-and-chain structures, and jellyfish microglia ^38^. Junqiu Jia et al. observed that during the recovery of WMI, microglia accumulate around the damaged myelin sheaths area and phagocytose myelin debris in the ischemic region in tMCAO mouse model, which display molecular characteristics associated with the CD11c+ microglial population ^39^. Moreover, the depletion of CD11c+ microglia exacerbates functional impairment in stroke mice and delays white matter regeneration ^39^. To unravel the molecular mechanism, our transcriptomic analysis pinpointed *Stap1* as a key downstream effector of IPA. According to the previous studies, STAP1 (Signal transducing adaptor protein 1) is a signal-transducing adaptor protein widely expressed in various cell types, including T cells and microglia ^40,41^. In T cells, STAP1 plays a crucial role by facilitating intercellular protein communication and signal transmission. T cells lacking STAP1 exhibit impaired signal transmission, reduced production of immune molecules, and are associated with the development of inflammation and autoimmune disease ^40^. However, the function of STAP1 in microglia has rarely been studied. Yang X et al. demonstrated that *Stap1* is involved in regulating neuroinflammation in microglia by influencing the cell’s activation status and phagocytic function ^42^. In this study, we had found that *Stap1* is expressed in microglia (Fig 6G); its expression is upregulated after ICH (Supplementary Fig 4) and further suppressed upon IPA supplementation. We therefore propose a novel mechanism wherein IPA alleviates the phagocytic inhibition exerted by *Stap1*, thereby ’releasing the brake’ on microglia and enabling more efficient debris clearance.

This study has obtained preliminary findings regarding the neuroprotective effect of the microbiota-gut-brain axis in patients with ICH. However, there is still room for improvement, and the study has several limitations as follows. One limitation of the present study is that IPA sensing and *Stap1* downregulation requires further validation. While our data position *Stap1* downstream of IPA, future studies are necessary to determine if this requires signaling through known receptors such as aryl hydrocarbon receptor (AhR) or Pregnane X receptor (PXR) ^43^ which could be tested using specific antagonists or genetic knockout models. The second limitation is that the mechanistic insight into microglia is limited. We could not ascertain the cellular specific of IPA versus other phagocytes, and we did not identify the specific microglial subtype responsive to IPA, and we relied on BV2 cell lines instead of primary microglia. Additionally, our data demonstrates that IPA could alleviated WMI and facilitate long-term motor function recovery, but we did not determine whether the preserved white matter integrity translates to sustained cognitive improvement, nor did it explore clinically viable strategies to deliver IPA to the brain. In summary, our findings provide a proof-concept that that gut metabolite IPA treatment enhances microglia phagocytosis through inhibit *Stap1* and WMI repair, offering a novel as a promising therapeutic strategy for ICH.

## Conclusions

The present study demonstrates that IPA could accelerate myelin debris clearance and promote neurological function recovery in mice after ICH. Since IPA is a gut microbiota derived metabolite, which is relatively safe and well-tolerated, our findings provide strong support for future clinical studies on intervention in gut microbiota (such as preparing probiotics or their metabolites) for the treatment of ICH.

## Article information

### Affiliations

Clinical Biobank Center, Microbiome Medicine Center, Guangdong Provincial Clinical Research Center for Laboratory Medicine, Department of Laboratory Medicine, Zhujiang Hospital, Southern Medical University, Guangzhou, 510280, China (M.P. M.Z., H.T., C.H., L.Z., Z.Z., Y.L., X.L., F.X., H.S.). The National Key Clinical Specialty, Neurosurgery Center, Engineering Technology Research Center of Education Ministry of China on Diagnosis and Treatment of Cerebrovascular Disease, Guangdong Provincial Key Laboratory on Brain Function Repair and Regeneration, The Neurosurgery Institute of Guangdong Province, Zhujiang Hospital Institute for Brain Science and Intelligence, Zhujiang Hospital, Southern Medical University, Guangzhou, 510280, China (M.P. M.Z., H.T., L.Z., Z.Z., H.S.). Department of Neurosurgery, Affiliated Hospital of North Sichuan Medical College, Nanchong, Sichuan Province, 637000, China (C.H., H.S.). Chinese Medicine Guangdong Laboratory State Key Laboratory of Traditional Chinese Medicine Syndrome, The Second Affiliated Hospital of Guangzhou University of Chinese Medicine, Guangzhou, Guangdong, China (S.L., X.C.). Guangdong Provincial Key Laboratory of Research on Emergency in TCM, Guangzhou 510120 Guangdong, China (S.L., X.C.).

### Author Contributions

H.S. conceived and supervised the study, acquired funding, oversaw project administration, and reviewed and edited the manuscript. M.P., M.Z and H.T. performed the experiments, analyzed the data and wrote the manuscript. C.H., L.Z. and Z.Z. contributed to the qPCR and behavior tests. Y.L contributed to the histology studies. X.L. and F.X. contributed to the UPLC-MS/MS. We thank S.L. and X.C for providing the valuable human cohort samples. The graphic abstract is created with by M.P. using BioRender. com, and we have been granted a license to use it. All authors approved the final article.

### Sources of Fundings

This research was funded by the National Natural Science Foundation of China (Grant No. 82571459), the Guangdong Provincial Clinical Research Center for Laboratory Medicine (Grant No. 2023B110008), and the Guangdong Basic and Applied Basic Research Foundation (Grant Nos. 2023A1515030045; 2025A1515010528).

### Disclosures

None.

### Footnote

Nonstandard Abbreviations and Acronyms.

## Notes

### Competing Interest Statement

The authors have declared no competing interest.

